# Homologous recombination contributes to the repair of acetaldehyde-induced DNA damages

**DOI:** 10.1101/2023.12.03.569808

**Authors:** Kosuke Yamazaki, Kazuto Takayasu, Trinh Thi To Ngo, Ayaka Onuki, Hideya Kawaji, Shunji Oshima, Tomomasa Kanda, Hisao Masai, Hiroyuki Sasanuma

## Abstract

Acetaldehyde, a chemical that can cause DNA damage and contribute to cancer, is prevalently present in our environment, e.g., in alcohol, tobacco, and food. Although aldehyde potentially promotes crosslinking reaction among biological substances including DNA, RNA, and proteins, it remains unclear what types of DNA damage are caused by acetaldehyde and how they are repaired. In this study, we examined acetaldehyde sensitivity of DNA damage-deficient cells established from human TK6 cell line. Among the mutants, mismatch repair mutants did not show a hypersensitivity to acetaldehyde, while cells deficient in base and nucleotide excision repair pathways increased its sensitivity. We found a delayed repair and hypersensitivity in homologous recombination (HR)-deficient cells but not in non-homologous end joining-deficient cells after acetaldehyde treatment. By analyzing the formation of acetaldehyde-induced RAD51 foci, which represent HR intermediates, HR-deficient cells, but not NHEJ, exhibits delayed repair of acetaldehyde-induced DNA damages, compared with *wild-type*. These results suggest that acetaldehyde causes complex DNA damages that requires various types of repair pathways. Interestingly, mutants deficient in TDP1 and TDP2, which are involved in the removal of protein adducts from DNA ends, exhibited hypersensitivity to acetaldehyde. the acetaldehyde sensitivity of the *TDP1^-/-^/RAD54^-/-^* double mutant was similar to that of each single mutant. This epistatic relationship between TDP1 and RAD54 suggests that that the removal of protein-DNA adducts generated by acetaldehyde needs to be removed for efficient repair by HR. Our study would help understand the molecular mechanism of genotoxic and mutagenic effects of acetaldehyde.

## Introduction

According to a report released by the World Health Organization (WHO), more than 3 million people suffers from various health problems due to harmful use of alcohol in 2016 [1]. Alcohol has been shown to be associated with cancer, such as oropharynx, oesophagus, liver, and breast cancers [1–3]. A cohort study in France by the International Agency for Research on Cancer strongly suggested that alcohol is the second leading risk factor for cancer, next to tobacco [4].

Alcohol dehydrogenase (ADH) oxidizes alcohol and generates acetaldehyde. Subsequently, it is metabolized by aldehyde dehydrogenase (ALDH), forming acetic acid [5]. The resultant acetic acid is decomposed into carbon dioxide and water in muscles and adipose tissues. In East Asia, there is a high frequency of genetic variants of the *ALDH2* gene showing a reduced activity in alcohol metabolism in these variant populations. The substitution of glutamic acid with lysine at amino acid 504 (Glu504Lys) in the *ALDH2* gene, which is seen in almost half of Japanese, exhibited symptoms of alcohol intolerance, such as nasal congestion, skin flushing, headaches due to the prolonged exposure to acetaldehyde [5,6]. The heterozygous genotype of an ALDH2 allele (*ALDH2*^+/Glu504Lys^) decreases alcohol metabolism activity to 1/16, compared with that of *ALDH2*^+/+^ genotype [7,8].

Aldehydes are produced in various metabolic reactions in cells. They can be present in certain foods like fruits and generated by oral bacteria [9]. Aldehydes, including acetaldehyde, has been demonstrated to have genotoxic potential. Aldehyde molecules non-enzymatically promote cross-linking reactions of molecules in close vicinity of DNA. Among them, the DNA-protein crosslinks (DPCs) are highly harmful for cell survival, because of their large size. DPCs become obstacles during DNA replication and transcription, subsequently leading to generation of DNA damage. DPCs are demonstrated to be removed by several repair pathways. Tyrosyl DNA phosphodiesterase proteins, TDP1 and TDP2, can remove DPCs enzymatically generated by topoisomerase 1 (TOP1) and 2 (TOP2) present at 3’ and 5’ ends of DNA respectively [10,11]. Recent studies demonstrated that TDP1 can remove not only TOP1 but also a variety of 3’ adducts from DNA for DNA repair [12]. DPCs can also be removed from genomic DNA in a nuclease-dependent manner. MRE11 nuclease, a factor involved in the repair of DNA double-strand break (DSB), removes DPCs together with the DNA strands via cleaving both DNA strands in the vicinity of the DPC [13–15]. Recent studies have demonstrated that *ALDH2*^-/-^/*FANCD2*^-/-^ mutant mice treated with ethanol exhibit symptoms similar to Fanconi anemia. This suggests that the Fanconi anemia pathway also plays an important role in repairing DPC caused by aldehydes. However, it remains unclear how DNA damages induced by aldehydes can be repaired.

In this study, we investigated the genotoxicity of acetaldehyde by analyzing acetaldehyde sensitivity of various DNA damage repair mutant cells. We show here that acetaldehyde generates complex DNA damages that require multiple repair pathways. We found that homologous recombination (HR) is one of the most important repair pathways for the repair of acetaldehyde-induced DNA damages, rather than the non-homologous end-joining (NHEJ) pathway. Acetaldehyde-induced DNA damages increased sister chromatid exchanges (SCEs) most likely due to the consequence of HR repair. The acetaldehyde-induced increase of SCEs was not observed in HR defective cells. This study will help understand the molecular mechanism underlying acetaldehyde-induced DNA damages and their repair.

## Materials and Methods

### Cell culture

Human TK6 B (TK6) cells were incubated in RPMI1640 medium (3026456, Nacalai Tesque, JP) supplemented with Horse serum (10%) (16050114, Gibco, US), penicillin (100 U/ml), streptomycin (100 µg/ml) (26253-84, Nacalai Tesque, JP), and Sodium pyruvate (200 mg/ml) (06977-34, Nacalai Tesque, JP)). Lenti-X^TM^ 293T cells (632180, TAKARA, JP) were maintained in DMEM supplemented with fetal bovine serum (10%) (Gibco, US), penicillin (100 U/ml), streptomycin (100 μg/ml), sodium pyruvate (200 mg/ml) and L-glutamine (16948-04, Nacalai Tesque, JP). TK6 and Lenti-X^TM^ 293T cells were maintained at 37_ under a humidified atmosphere and CO_2_ (5%).

### Lentivirus production and isolation of Azami-green-positive TK6 cells

Lentiviral vectors, which can express Azami green, were simultaneously transfected with virus-packaging plasmids into LentiX-293T cells. Lentiviruses were harvested at 48 h post-transfection. TK6 cells were infected with the virus for 48 h and then Azami-green (AG)-positive cells were selected by cell sorter, Aria III (Becton, Dickinson and Company, US). The selected cells were seeded into 96-well plates to isolate single clones. The AG-positive clones were stocked after the minimal passage to avoid reduction of green-fluorescence signal due to lentivirus element-derived silencing in TK6 cells.

### Sensitivity assay using FACS

AG-positive and DNA repair mutant cells were incubated in the logarithmic phase. After counting the cell density by Countess2 (AMQAX100, Invitrogen, US), an equal number of cells were mixed. The mixed cells were exposed to the indicated concentration of acetaldehyde diluted in PBS for 30 min, and then washed with the normal medium. To incubate the cells for five days, we diluted the cells two or three times every day to avoid 100 % confluent.

### Methylcellulose cell survival assay

The number of TK6 cells treated with acetaldehyde was accurately counted by Countess2. The cells were incubated in the medium containing methylcellulose (1.6%) (M0387, SIGMA, US), NaHCO2 (0.4%) (31212-25, Nacalai tesque, JP), 1 x DMEM/HAM (D8900, SIGMA, US), Horse serum (10%), and Sodium pyruvate (1mM). The number of colonies was counted 10-12 days after the cell seeding.

### Immunostaining and microscopic analysis

Cells were treated with acetaldehyde and subsequently washed twice with cold PBS to remove acetaldehyde completely from the medium. Cells were collected using a cytospin (Thermo Fisher Scientific, US) and subjected to fixation by formaldehyde (4%) (061-00416, Fujifilm, JP) diluted by PBS, permeabilization by Tween-20 (0.1%) (23926-35, Nacalai Tesque,JP) in PBS, and blocking by BSA (5%) (01683-48, Nacalai Tesque,JP) in PBS. The 1st and 2nd antibodies were anti-RAD51 (70-001, 1/1000 dilution, Bioacademia, JP) and goat anti-rabbit conjugated with alexa488 (A-11001, 1/1000 dilution, Invitrogen, US). The slides were mounted in Fluoro-KEEPER containing 4’, 6-diamidino-2-phenylindole (DAPI) (12745-74, Nacalai Tesque, JP). The images were taken by SP8 and STELLARIS confocal microscope (Leica, DE).

### RNA-seq

Total RNA was extracted by the RNeasy Mini kit from QIAGEN (74104, Hilden, DE). The library preparation and sequencing were performed by Eurofins Genomics (Tokyo, JP). The RNA-seq reads were processed by fastp (version 0.23.2) to remove poor-quality reads and adaptor sequences [16]. The resultant fastq files were mapped by STAR (version 2.5.3) along the reference genome (GRCh38) [17,18]. The expression levels of each gene were calculated by RSEM (version 1.3.3) as transcripts per million (TPM) [19]. The gene expression of the indicated genes was analyzed by Seaborn (version 0.12.2) for data visualization.

### Chromosome analysis

1 x 10^6^ cells were treated with colcemid (Final conc. 0.1 μg/mL)(15212012,Thermo Fisher Scientific, US) for 3 h. Cells were suspended in potassium chloride (KCl, 75 mM) for 15 min. Carnoy’s solution (a 3:1 mixture of methanol and acetic acid) was added into cell suspension with KCl (28513-85, Nacalai tesque, JP) solution and mixed gently. After the centrifuge (1000 rpm, 5min), the cell was suspended in 500 µL and dropped on slides. The slides were dried on the plate warmed at 55 °C and stained with a Giemsa solution (5%) (079-04391, Fujifilm, JP) for 10 min. The chromosome breaks were detected by KEYENCE BZ-700 (Tokyo, JP) with x100 lens.

### Sister chromatid exchange analysis

0.8 x 10^6^ cells in 1 mL were incubated with BrdU (final conc. 20 µg/mL) (B1575, Tokyo Kasei, JP) for 28 h. The cells were treated with colcemid (0.1 μg/mL) for 3 h. The cells were treated with 1mL of 75 mM KCl, left on a bench for 20 min, and then incubated with 5 mL Carnoy’s solution for 30 min. The cells were washed twice by 5 mL Carnoy’s solution. The slides were washed with 50 % ethanol and then dried on a hotplate heated at 42 °C. The cell suspension was dropped on the warmed slide. The slides were incubated with 50 mM phosphate buffer (pH 6.8) (31801-05 and 31726-05, nacalai tesque, JP) containing Hoechst 33258 (H3569, Invitrogen, US) at 4 °C for 20 min. The slides were washed with McIlvaine solution (164 mM Na2HPO4 and 16 mM citric acid (pH 7.0)) for 20 min. The slides were covered with a coverslip to avoid drying and exposed to blacklight (365 nm UV) for 1 h. The slide was incubated with the following reagents: McIlvaine solution for 1 min, 2 X SSC for 62 °C for 1h, 50 mM phosphate buffer (pH 6.8). The chromosomes were visualized by staining 3% Giemsa solution diluted in 50 mM phosphate buffer (pH 6.8) for 30-60 mins.

### Quantification and Statistical Analysis

For all statistical analyses with a *p-value*, Mann-Whitney U test was used. Error bars represent standard deviation (SD), as indicated in the legends. The statistical analysis was performed using PRISM 8 (version 8.4.2).

## Results

### Establishment of a method for measuring sensitivity to drugs and ionizing radiation

To establish the conventional method for drug sensitivity, we have generated lenti-virus expressing Azami-green (AG) fluorescent protein and infected it into *wild-type* TK6 cells [20]. After the infection, we subcloned cells expressing AG. The expression of AG did not affect cell proliferation, compared with *wild-type* cells. We evenly mixed *wild-type* cells expressing AG (AG-positive cells) and *RAD54*^-/-^ cells (AG-negative cells) (Figure 1A), which has shown hypersensitivity to IR (Figure 1B) [21]. We then examined the ratio of AG-positive and negative cells after treatment of ionizing radiation (IR) by fluorescence-activated cell sorting (FACS). If the mutant deficient in the repair of IR-Induced damages dies or grows slowly after IR treatment, we expect that AG-positive and -negative ratio would be changed, compared with that of untreated cells. Indeed, IR treatment reduced AG-negative/AG-positive ratio in *RAD54*^-/-^ and *LIG4* ^-/-^ cells (Figure 1C). We therefore conclude that the novel assay using FACS can indicate sensitivity to X-rays (Figure 1C). Compared with the colony formation assay using methylcellulose, the established methods here, named “AG assay”, is less laborious and timesaving for checking drug sensitivity.

**Figure 1.**
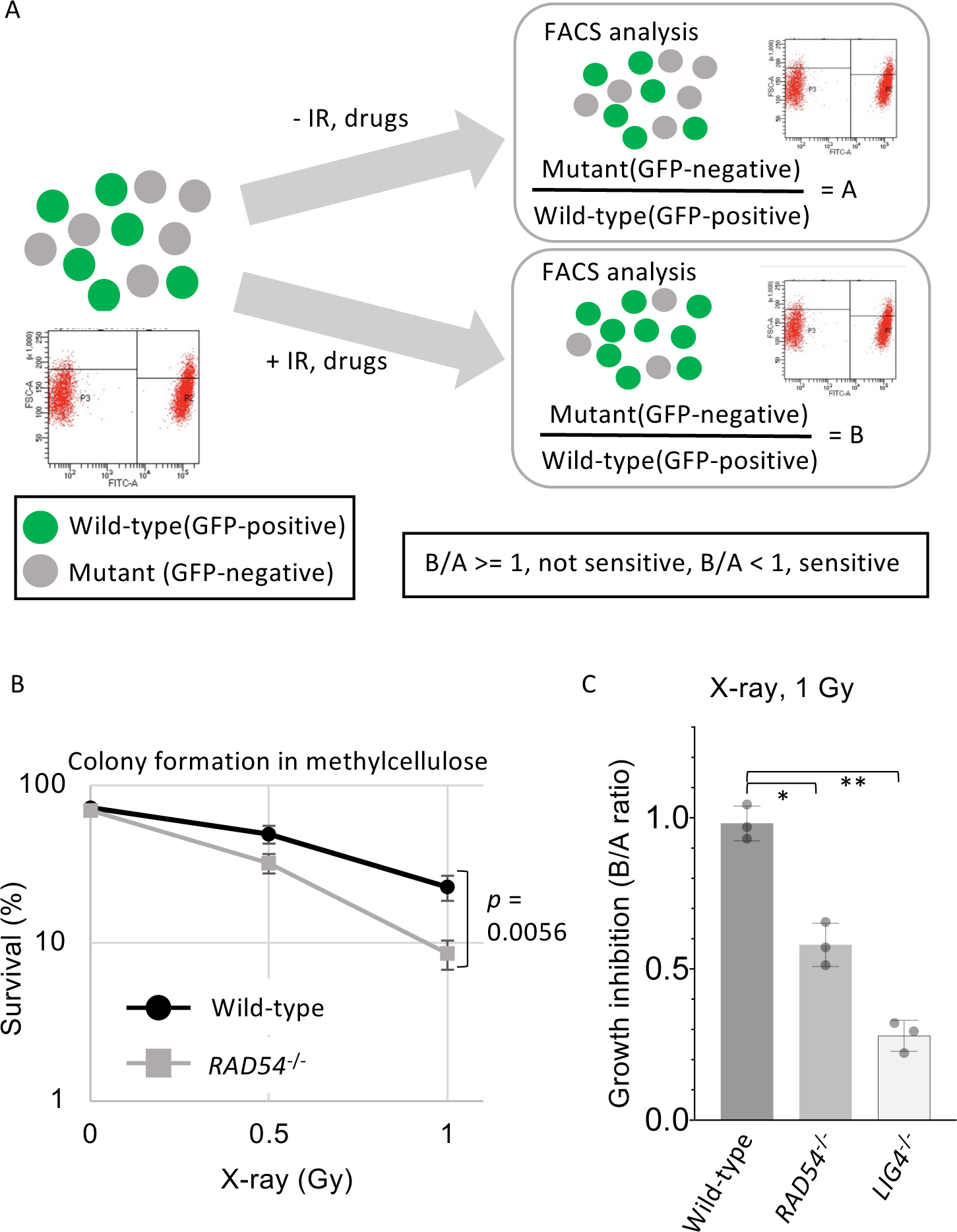
Schematics of flow cytometry-based sensitivity assay. The experiments were performed three times independently. Asterisks and error bars indicate *p-value* < 0.05 calculated from student t -test and standard deviation (SD) (A) Experimental scheme of sensitivity assay using flow cytometry (FACS). (B) X-ray sensitivity to *wild-type* and *RAD54^-/-^* TK6 cells. After exposure of the cells to the indicated dose, the cells were seeded in a methylcellulose medium. The cells were incubated for eight days and then the colonies were counted. (C) Sensitivity assay using the method described in (A).

### Optimizing the experimental condition for acetaldehyde sensitivity assay

Several reports demonstrated acetaldehyde genotoxicity in vivo [22]. Biochemical analysis suggests that acetaldehyde promotes interstrand crosslinks in DNA [23]. We sought to examine acetaldehyde genotoxicity using DNA-repair deficient TK6 mutant cells [13,21,24,25]. We initially exposed the cells to acetaldehyde by incubating them in the acetaldehyde-containing medium at 37 °C. We were not able to find significant sensitivity to acetaldehyde of DNA repair mutants. Since the boiling temperature of acetaldehyde at 101.3 kilopascals is approximately 20.2 °C. This is most likely due to the evaporation of acetaldehyde in the 37 °C incubator. To prevent the volatilization and detoxification in the medium, we suspended the cells in PBS and then treated them with 100 µM acetaldehyde in 1.5 mL tube at room temperature for 30 min. We then incubated the cells in the medium containing serum. To optimize the concentration of acetaldehyde to measure cell survival, we transiently treated the *wild-type* and *RAD54*^-/-^ cells with different concentrations of acetaldehyde for 30 min. We detected hypersensitivity to acetaldehyde in *RAD54*^-/-^ cells with methylcellulose assay (Figure 2A) and AG assay (Figure 2B).

**Figure 2.**
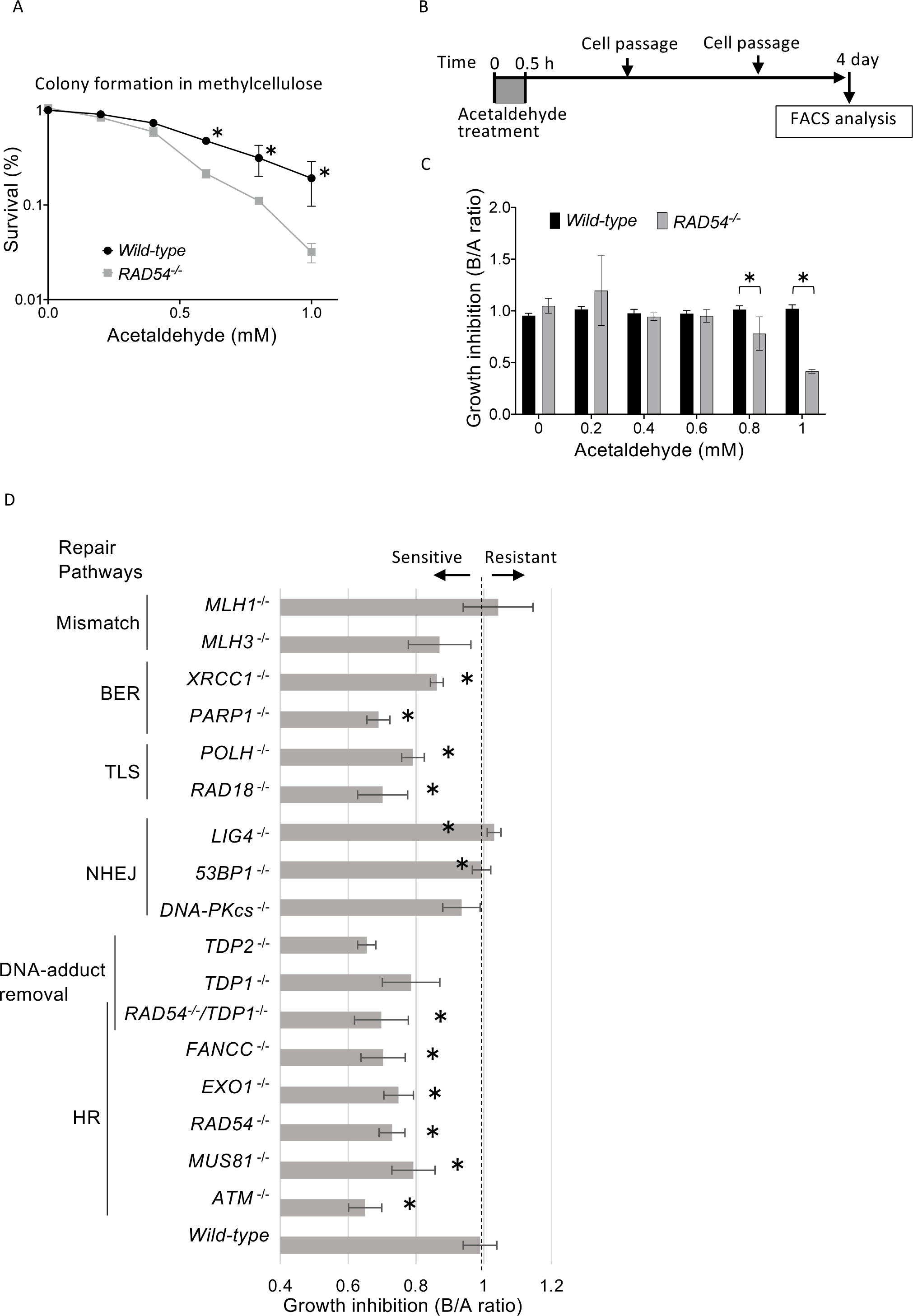
Cell sensitivity assay to acetaldehyde using FACS. The experiments were performed three times independently. Asterisks and error bars indicate *p-value* < 0.05 calculated from student t -test and standard deviation (SD) (A) Acetaldehyde sensitivity using methylcellulose medium. After the exposure to acetaldehyde in 1.5 mL test tube, for 30 min, the cells were seeded in a methylcellulose medium. The cells were incubated for eight days and then the colonies were counted. (B) Experimental design to analyze the acetaldehyde sensitivity at the indicated concentration. The cells were exposed to the indicated dose of acetaldehyde for 60 min in 1.5 mL test tube, and then washed with the medium. The cells were diluted several times by cell passage and incubated for four days. The sensitivity was measured by FACS and calculated by the formula described in Figure 1A. (D) Result of Sensitivity assay to acetaldehyde. The experimental procedure was followed by (B).

**Figure 3.**
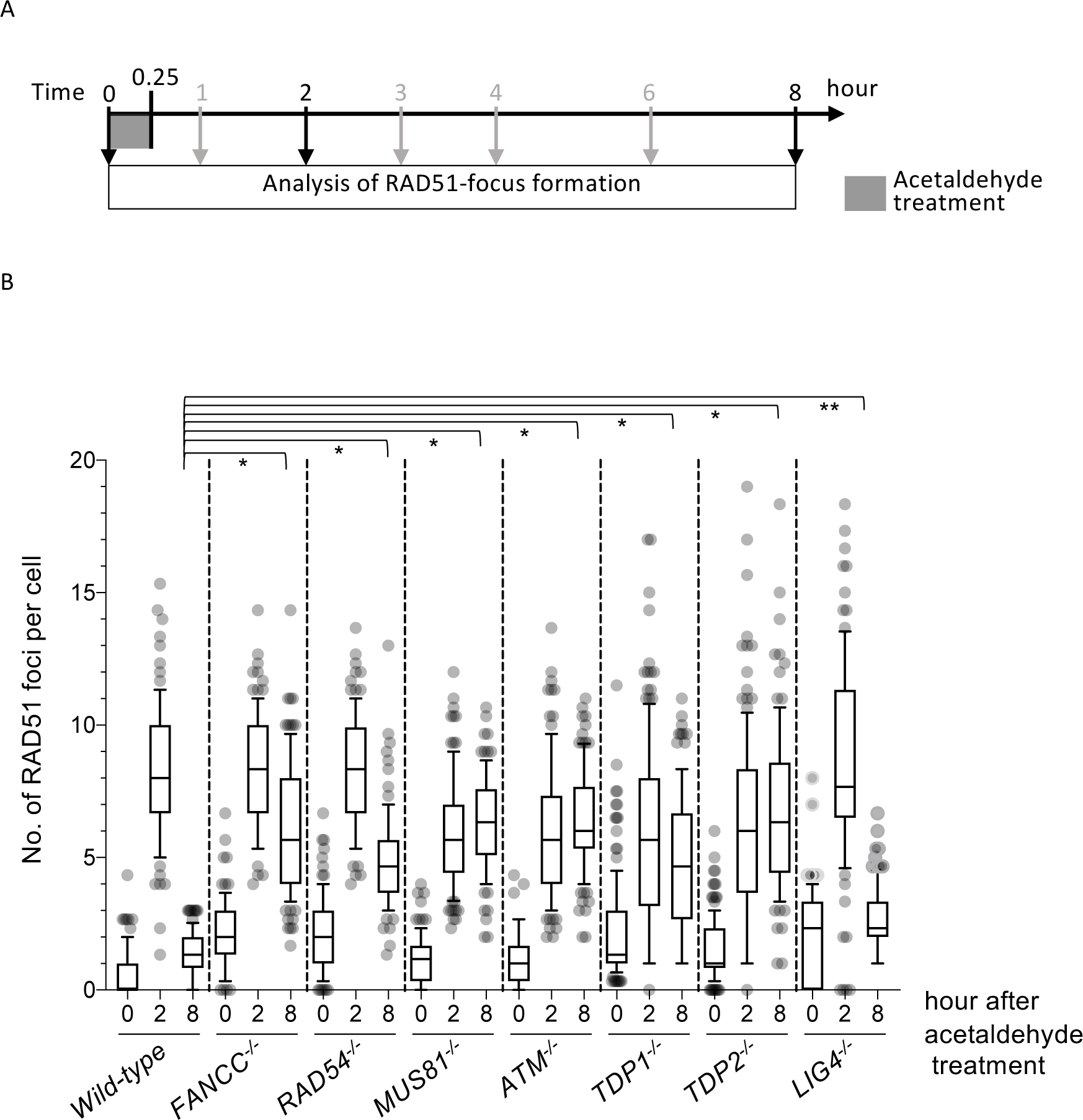
RAD51-focus formation in response to acetaldehyde exposure. The cells were initially exposed to 100 µM acetaldehyde for 15 min in a 1.5 mL test tube and then started incubating the cells in the normal medium (time 0.25). The samples were taken at the indicated time-points. The cells were stained with anti-RAD51 antibody. The experiments were performed three times and calculated *p-values*. The single asterisks indicate *p-value* < 0.05 calculated from student t -test. Double asterisk indicates *p-value* > 0.05. Bounds of box are 25–75th percentile, the center shows the median, whiskers indicate the 10–90 percentiles, data points outside this range are drawn as individual dots.

### Sensitivity assay to acetaldehyde using DNA repair-deficient mutant cells

To examine the genotoxicity of acetaldehyde, we exposed a variety of DNA repair-deficient mutant cells to acetaldehyde. Mismatch repair-deficient cells, *MLH1*^-/-^ and *MLH3*^-/-^ [25], did not show sensitivity to acetaldehyde (Figure 2B), indicating that the mismatch repair pathway does not play an important role in the repair of acetaldehyde-induced damages. In contrast, we observed a significant reduction in AG ratio in *XRCC1*^-/-^ and *PARP1*^-/-^ cells after exposure to acetaldehyde. Since XRCC1-PARP1 is involved in the base-excision repair (BER) pathway to promote the repair of damaged bases [26], this result indicates that BER pathway contributes to repairing acetaldehyde-induced damages. Interestingly, mutant cells deficient in homologous recombination (HR), *FANCC*, *EXO1*, *RAD54*, *MUS81*, and *ATM*, all exhibited hypersensitivity to acetaldehyde, indicating a critical role of HR in DNA damage repair induced by acetaldehyde [25]. In contrast, *LIG4*, mutants deficient in *DNA-PKcs*, and *53BP1*, genes involved in non-homologous end joining (NHEJ) pathway, were not sensitive to acetaldehyde [24,27]. These results indicate that acetaldehyde-induced DNA damages are repaired by homologous recombination, rather than non-homologous end joining.

The mutants in tyrosine-phosphodiesterase 1 (TDP1) and 2 (TDP2), which are enzymes to remove topoisomerase1 (TOP1) adduct and topoisomerase2 (TOP2) adduct, respectively, covalently bound to DNA ends, exhibited sensitivity to acetaldehyde higher than in *wild-type* cells [13]. Several studies demonstrated that TDP1 can remove not only TOP1 adduct but also a wide range of 3’ synthetic DNA adducts as a substrate. The result suggests a significant contribution of TDP1 and TDP2 to the repair of bulky DPCs induced by acetaldehyde.

We also found acetaldehyde hypersensitivity in the cells deficient in translesion synthesis (TLS) and template-switch (TS) pathways, *POLH*^-/-^ and *RAD18*^-/-^ cells [28,29]. Considering the idea that acetaldehyde causes bulky chemical adducts along genomic DNA, these results suggest that TLS and TS pathways contribute to bypassing bulky adducts generated by acetaldehyde during DNA replication.

### Homologous recombination is required for efficient repair of acetaldehyde-induced DNA damages

To examine the contribution of HR to the repair of acetaldehyde-induced DNA damages, we examined RAD51-focus formation upon transient exposure to acetaldehyde. In the *wild-type* cells, the number of RAD51 foci was increased at 2 h after the addition of acetaldehyde and then decreased at 8 h. Consistent with acetaldehyde sensitivity (Figure 2C), we found that the repair kinetics was delayed in *RAD54*, *MUS81*, *ATM*-deficient cells. The delayed repair was also seen in the mutants deficient in *FANCC* and *TDP1* genes. Consistent with the previous findings [30,31], these results indicate that Fanconi anemia and DPC repair pathways contribute to the repair of acetaldehyde-induced DNA damage. To examine the genetic interaction between genes involved in DPC removal and HR pathways, we generated *TDP1*^-/-^/*RAD54*^-/-^ double mutant cells. Acetaldehyde sensitivity in *TDP1*^-/-^/*RAD54*^-/-^ double mutant cells was similar to that in *RAD54*^-/-^ single mutant cells, indicating an epistatic relationship. HR is likely necessary for DSB repair after removing the bulky adducts covalently associated with DNA ends by TDP2.

### RNA expression profiles of ethanol metabolism-related genes in TK6 cells

We sought to examine the potential activity of alcohol metabolism in TK6 cells. We extracted RNA from TK6 cells without any treatment and then analyzed RNA expression by next-generation sequencing. We detected the expression of DNA damage repair-related genes, including *RAD54*, *MUS81*, *FANCC*, *ATM*, *TDP1*, and *LIG4* (Figure 4A). The primary enzymes related to alcohol metabolism are alcohol dehydrogenase (ADH) and aldehyde dehydrogenase (ALDH). Multiple ADH and ALDH enzymes are encoded on different alleles. RNA-seq profiles showed a significant expression of ADH5. Several studies demonstrated that ADH5 is involved in ethanol metabolism [32,33]. We also detected significant expression of ALDH1B, which is known to be localized in mitochondria like ALDH2 and is involved in acetaldehyde detoxification [34]. TK6 cells, therefore, would have the potential to metabolize ethanol.

**Figure 4.**
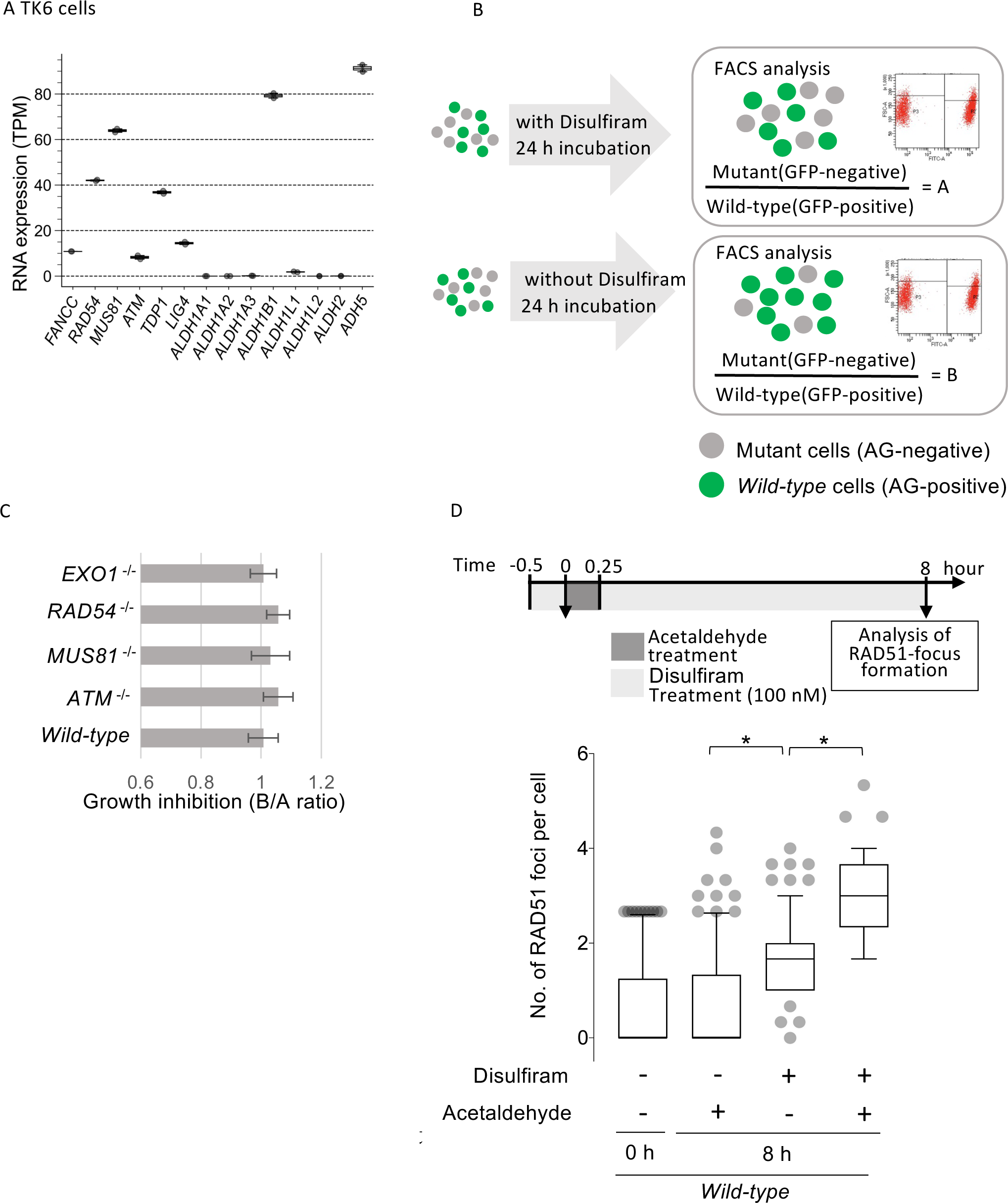
Simultaneous treatment of disulfiram and acetaldehyde increases RAD51 foci. (A) RNA expression profiles of DNA repair-related and alcohol and aldehyde-related genes in TK6 cells. The total RNA was extracted from logarithmic-phase cells of *wild-type* without any treatment. The read counts of individual transcripts were shown as transcripts per million (TPM). The RNA-seq analysis was independently performed twice. (B)(C) Experimental design to examine the effect of disulfiram on cell growth in TK6 cells. After mixing *wild-type* (AG-positive) and mutant (AG-negative) cells, they were split and incubated for 24 h. GFP-positive and -negative ratios in disulfiram-treated (upper) and -untreated (lower) were calculated as value “A” and value “B”, respectively. Growth defects by disulfiram were calculated by B/A ratio as shown in (C). The experiments were performed three times independently. (D) Effect of disulfiram on RAD51 focus formation. The upper diagram represents the experimental scheme. Disulfiram was added 30 min before acetaldehyde treatment to inactivate the enzymes completely. The experiments were performed three times independently.

### Effect of disulfiram on the repair of acetaldehyde-induced DNA damages in TK6 cells

To examine the activity of acetaldehyde oxidation in TK6 cells, we treated *wild-type* cells with acetaldehyde in the presence of tetraethylthiuram disulfide, disulfiram, an inhibitor of ALDH2 and ALDH1 enzyme. Disulfiram is a well-known anti-alcohol agent that inhibits ALDH, resulting in the accumulation of acetaldehyde and extreme ‘hangover’ symptoms. We firstly exposed the cells with disulfiram for 8 h and then mixed them with *wild-type* cells expressing Azami Green (Figure 4B). After 24 h incubation with a normal medium, we measured AG+/-ratio by FACS analysis as was conducted in Figure 1. We analyze several HR-deficient mutants. We did not detect growth inhibition in the medium containing 100 nM disulfiram (Figure 4C), indicating that disulfiram does not have cytotoxicity at least in this condition. Under the same condition, we transiently exposed the cells to acetaldehyde (Figure 4D) and then analyzed RAD51-focus formation. The number of RAD51 foci at 8 h in *wild-type* cells was comparable to that at 0 h in *wild-type* cells. In contrast, when we incubated the cells with disulfiram, the number of RAD51 foci significantly increased. Inhibition of the metabolism of endogenous aldehyde species possibly causes DNA damage that can be repaired by RAD51. Acetaldehyde treatment together with disulfiram increases the number of RAD51 foci compared with that in cells treated with disulfiram alone. Aldehyde species are constantly generated by metabolism in the cell, and the inhibition of ALDH2 and ALDH1 activities by disulfiram causes an increase in the intracellular concentration of aldehyde species, resulting in the accumulation of DNA damages.

### Acetaldehyde increases the number of chromosomal breaks and sister chromatid exchange

We examined acetaldehyde genotoxicity by chromosome analysis. We treated the cells with acetaldehyde for 1 h and incubated them for another 10 h (Figure 5A). We added colcemid to enrich metaphase cells 3 h before cell harvest. In *wild-type* cells, the number of breaks in acetaldehyde-treated cells did increase compared to untreated cells (Figure 5B), indicating unrepaired acetaldehyde-induced DNA damages could be detected as chromosome breaks. We next examined the effect of acetaldehyde on sister chromatid exchange (SCE). A number of studies demonstrated that HR-deficient mutants decreased SCE frequency [35–38]. If the HR pathway contributes to the repair of acetaldehyde-induced repair, we anticipated that HR deficiency suppresses SCE frequency. We therefore analyzed HR-deficient *RAD54* mutant cells. *BLM*-deficient cells have been demonstrated to exhibit an increase of spontaneous SCE[39]. Acetaldehyde treatment increased the number of SCEs compared with untreated *wild-type.* This increased number was suppressed by RAD54 loss, suggesting that HR causes an increased number of SCEs. These results indicate that acetaldehyde-induced DNA damage is repaired by HR.

**Figure 5.**
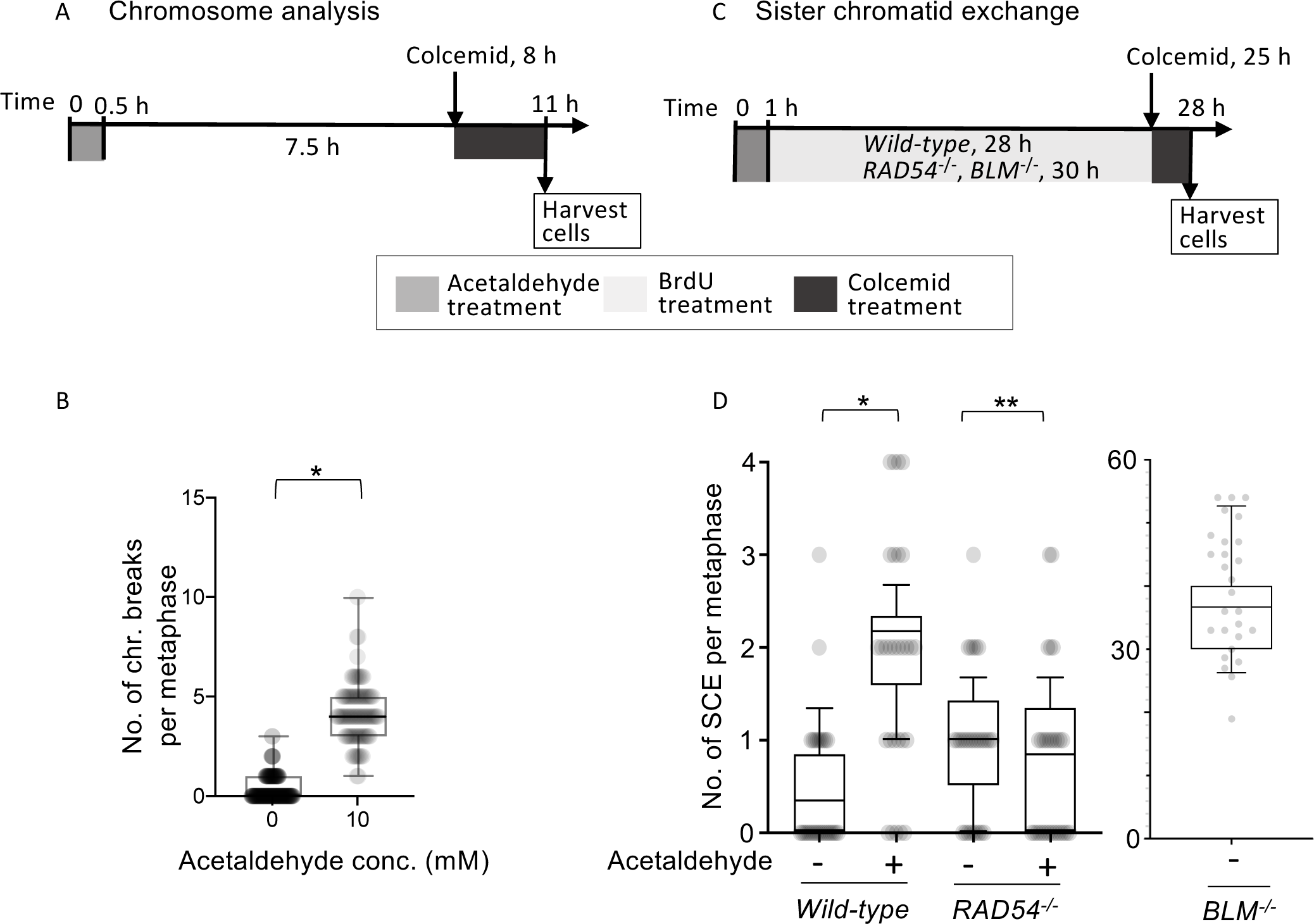
Increased aberrant chromosomes in response to acetaldehyde. (A) Experimental scheme of chromosome analysis. (B) The *wild-type* cells were treated with 100 µM acetaldehyde for 30 min in PBS and then incubated with the normal medium for 4.5 hours. Celcemid was added for the enrichment of mitotic cells. The bounds of the box are 25–75th percentile, the center shows the median, and the whiskers indicate the 10–90 percentiles. 50 metaphase cells were counted for each experiment. (C) Experimental scheme of sister chromatid exchange analysis. The *wild-type* and mutant (*RAD54*^-/-^ and *BLM*^-/-^) cells were incubated with BrdU for 28 and 30 hours, respectively, relevant to two rounds of the cell cycle. The asterisk indicates *p-value* < 0.05, calculated by student-t-test. (D) The single and double asterisks indicate *p-values* < and > 0.05, respectively. The bounds of the box are 25–75th percentile, the center shows the median, and the whiskers indicate the 10–90 percentiles. The experiments were independently performed three times. 10 metaphase cells were counted for each experiment.

## Discussion

This study revealed that acetaldehyde induces complex damage requiring various repair pathways. Among the acetaldehyde-sensitive mutant cells, TDP1 and TDP2 are enzymes involved in the removal of DNA-protein crosslinks (DPCs). These results suggest the potential involvement of these factors in removing protein adducts crosslinked to DNA by acetaldehyde. It is known that TDP1 and TDP2 do not have efficient capabilities for intact TOP1 and TOP2 covalently attached to DNA ends, respectively. TOP1/2 covalently bound to DNA needs to be degraded to the peptide level for subsequent removal by TDP1/2. Therefore, it is possible that proteasome and/or proteases need to firstly digest protein crosslinked to DNA to reduce the size of protein adducts caused by acetaldehyde. So far, SPRTN, a homolog of Wss1 in yeast, is a candidate protease for the process. In TK6 cells, since *SPRTN*^-/-^ cells we have generated resulted in extremely poor proliferation [40], we were unable to measure acetaldehyde sensitivity. Previous studies or findings suggest that after SPRTN or the proteasome promotes the degradation of bulky protein adducts into peptides, endonuclease(s) introduce nicks to remove DNA together with the attached peptides. Further investigation is needed to identify the molecular mechanism of how endonucleases are recruited onto the peptide-conjugated sites. In particular, during the S phase, when DNA replication forks encounter bulky protein adducts, the replication process is halted, resulting in the exposure of single-stranded DNA regions containing bulky protein adducts. It is known that, during DNA replication, the Poly ADP-Ribosylation of PARP1 and XRCC1-dependent repair pathways is involved in the maturation of Okazaki fragments by promoting the recruitment of FEN1 nuclease [41]. Even in the case of exposure of single-stranded DNA regions containing bulky protein-DNA adducts in S phase, it is plausible that the PARP1 and XRCC1 promote the recruitment of FEN1 and MRE11 nucleases as they do for Okazaki fragment maturation. The acetaldehyde sensitivity observed in *XRCC1* or *PARP1*-deficient cells may be due to the defects in the recruitment of nucleases, such as FEN1 and MRE11 nucleases. The mutants in TLS and TS pathways also exhibited hypersensitivity to acetaldehyde. Although the mechanism of the TS pathway is poorly understood, TLS pathway is involved in bypassing the unhooked cross-linked nucleotides, and thus restores the nascent strand. It would contribute to bypassing the bulky protein-DNA adducts generated by acetaldehyde during DNA replication.

Ingestion of 0.4g ethanol per kilogram of body weight, which is roughly equivalent to a 60 kg person consuming 500 mL beer (Alcohol, 5 %), increases acetaldehyde by approximately 80 µM in peripheral blood of individuals with the ALDH2^Glu504Lys^ homozygous variant [42]. Additionally, other studies demonstrated that short-term alcoholic beverage exposure in the mouth increases acetaldehyde concentration up to 1 mM in saliva [43,44], suggesting that acetaldehyde would be produced in a different concentration dependent on tissue and/or tissues. Our study has successfully detected the potential DNA toxicity of acetaldehyde at a much lower acetaldehyde concentration than that in saliva by using DNA repair mutant cells. We have used 100 µM of acetaldehyde, which is considered to be mimic the physiological events occurring when hematopoietic cells are exposed during alcohol metabolism [45]. To prevent hazardous and harmful drinking, it is crucial to provide individuals with scientific insights into the toxicity of alcohol. This study offers valuable scientific information that can support these educational efforts.

## Acknowledgement

We would like to thank the members of Genome Dynamics laboratory in TMiMS for their helpful comments and suggestions on the manuscript. Computations were partially performed on the NIG supercomputer at ROIS National institute of Genetics. This work was conducted through the research grant from Asahi Quality and Innovations and the Ministry of Education, Science, Sport and Culture to H.S (KAKENHI 19H04267).

## AUTHOR CONTRIBUTIONS

This study was conceived by H.S. Experiments and data analysis were performed by K.Y., K.T., T.T.T.N, A.O., H.K., S.O., T.K. H.M., and H.S. The paper was written by H.S.

---

**Figure S1.**
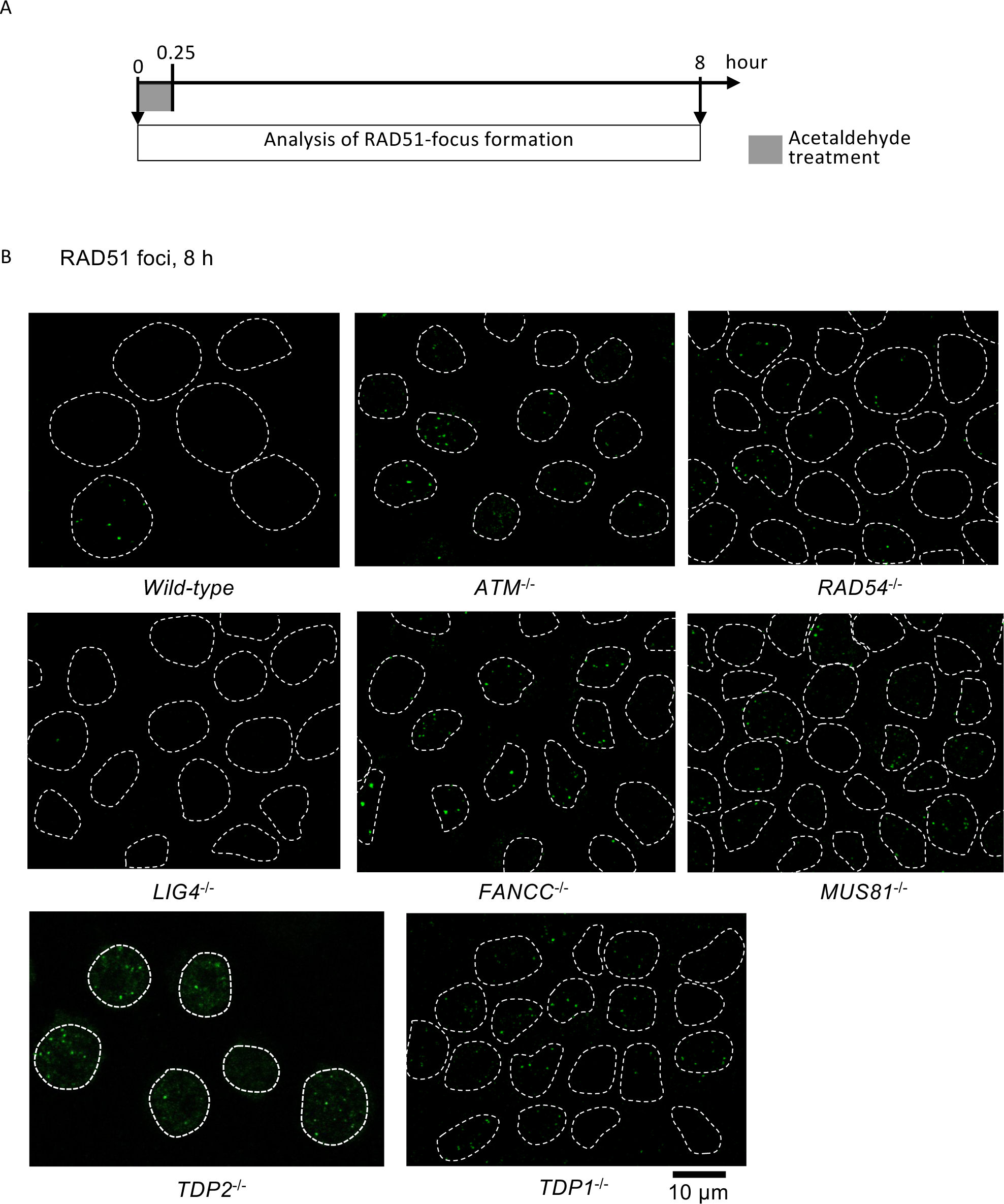
RAD51 focus formation in response to acetaldehyde exposure. Related to Figure 3. The cells were initially exposed to 100 µM acetaldehyde for 15 min in a 1.5 mL test tube and then started incubating the cells in the normal medium (time 0.25). (A) The samples were taken at the indicated time-points. (B) representative images of RAD51 foci at eight hours.

**Figure S2.**
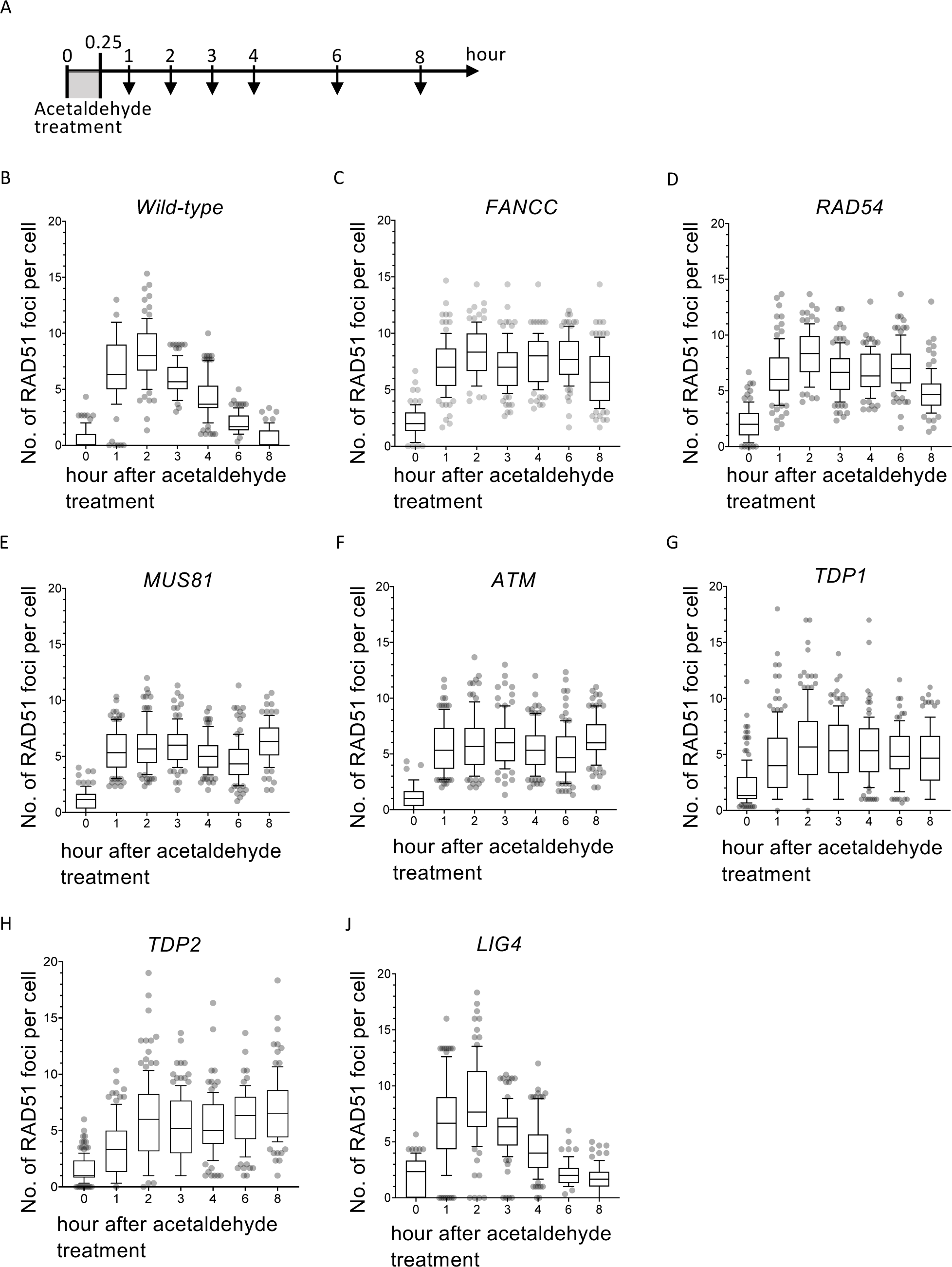
RAD51 focus formation in response to acetaldehyde exposure. Related to Figure 3. The cells were initially exposed to 100 µM acetaldehyde for 15 min in a 1.5 mL test tube and then started incubating the cells in the normal medium (time 0.25). (A) The samples were taken at the indicated time points. (B) The kinetics of RAD51 focus formation in response to acetaldehyde at the indicated time points. Figure 3 shows 0, 2, and 8 hours of the indicated cells.

**Figure S3.**
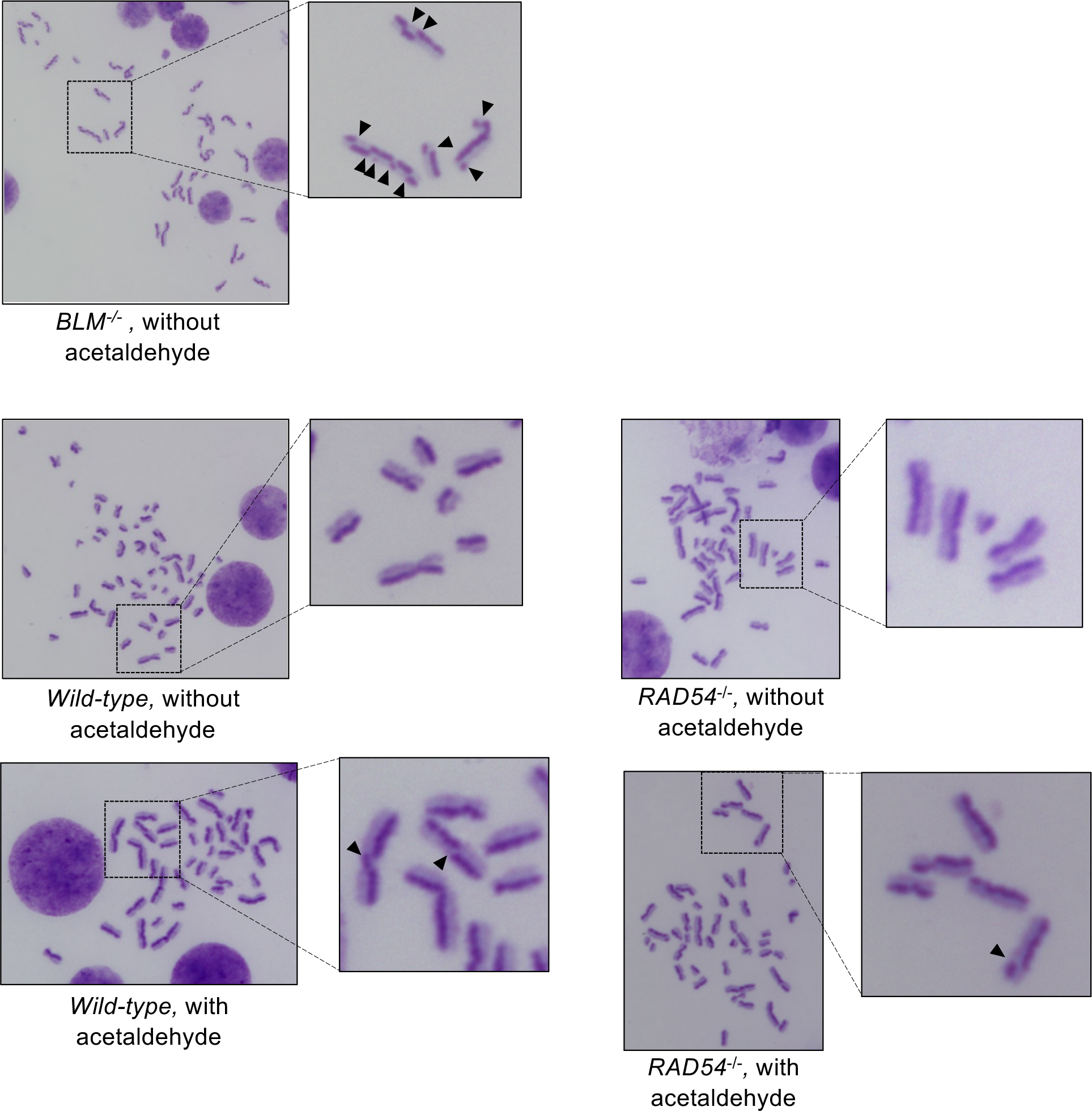
Representative images of SCE. Related to Figure 5. The cells were exposed to 100 µM acetaldehyde for 15 min in a 1.5 mL test tube and then started incubating as described in Figure 5C. Arrowheads indicate exchange points between sister chromatids.

**Figure S1 B.**
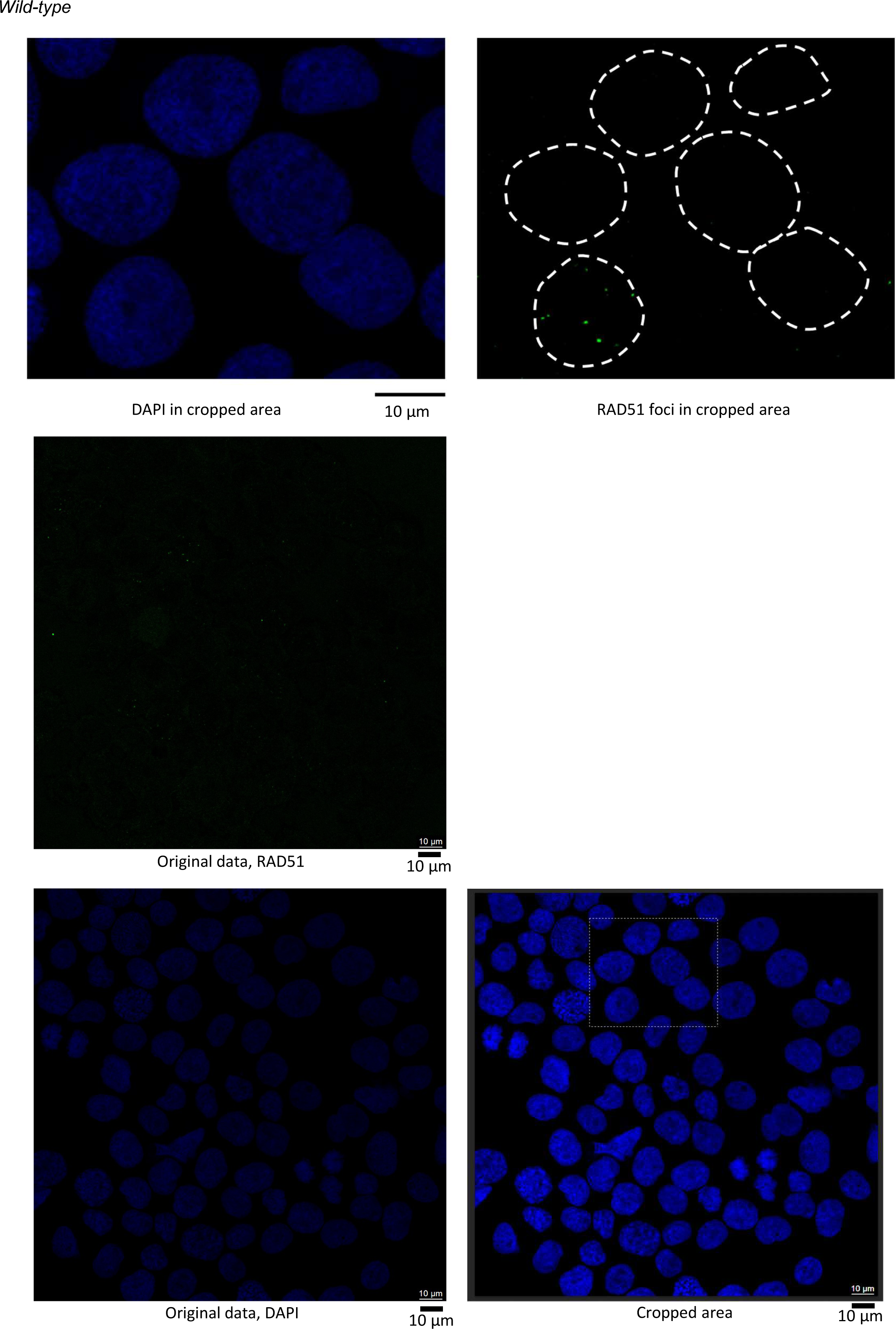
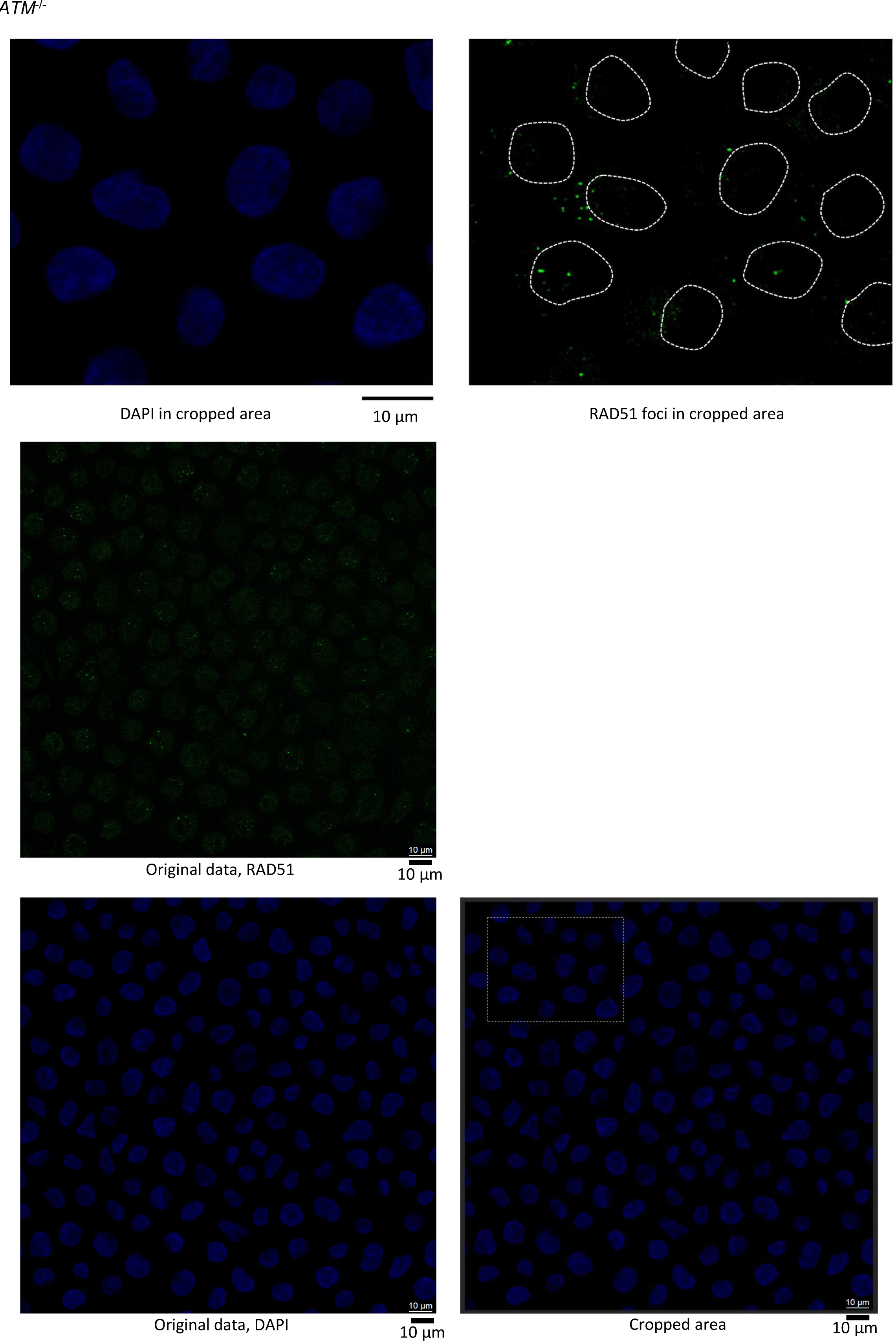
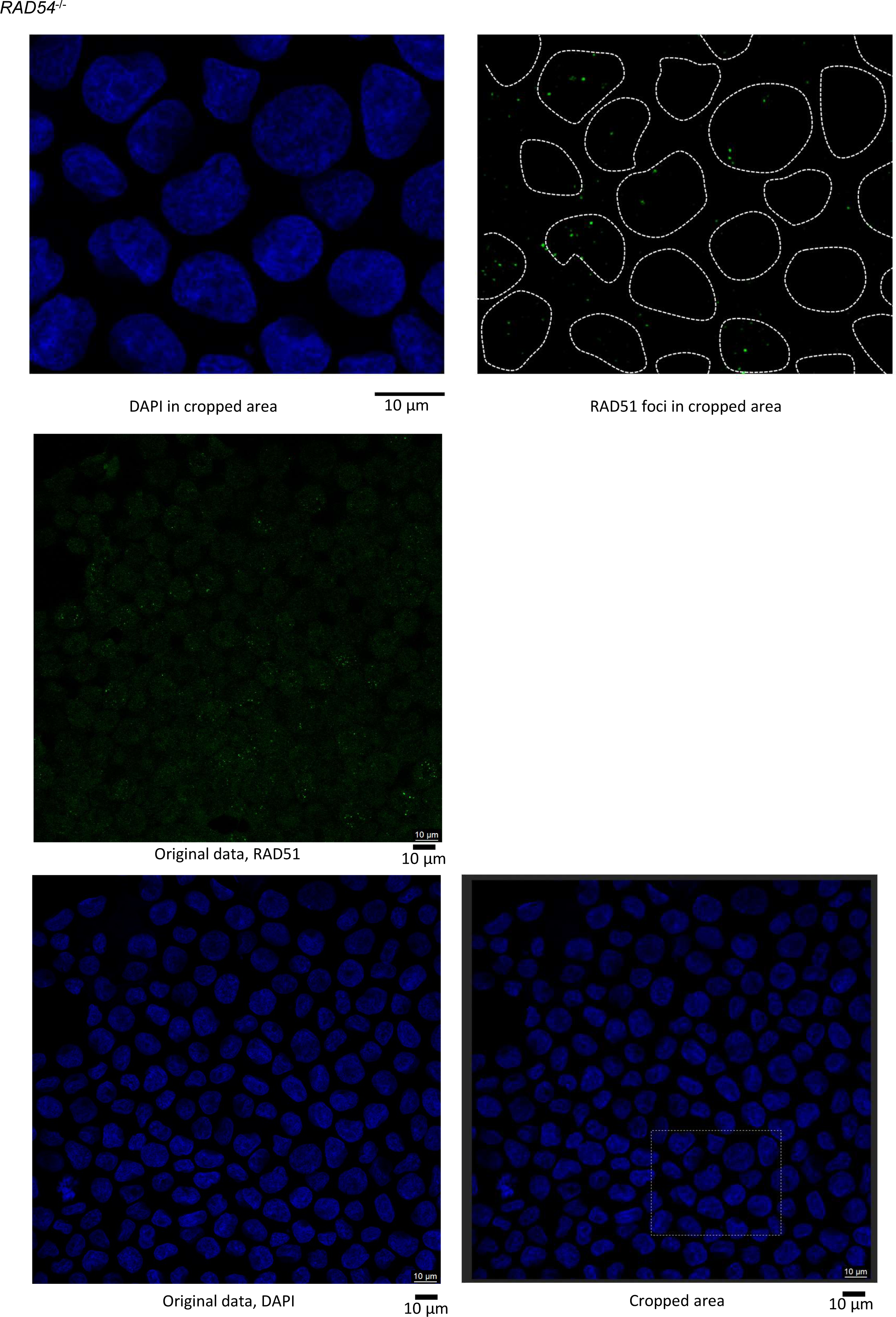
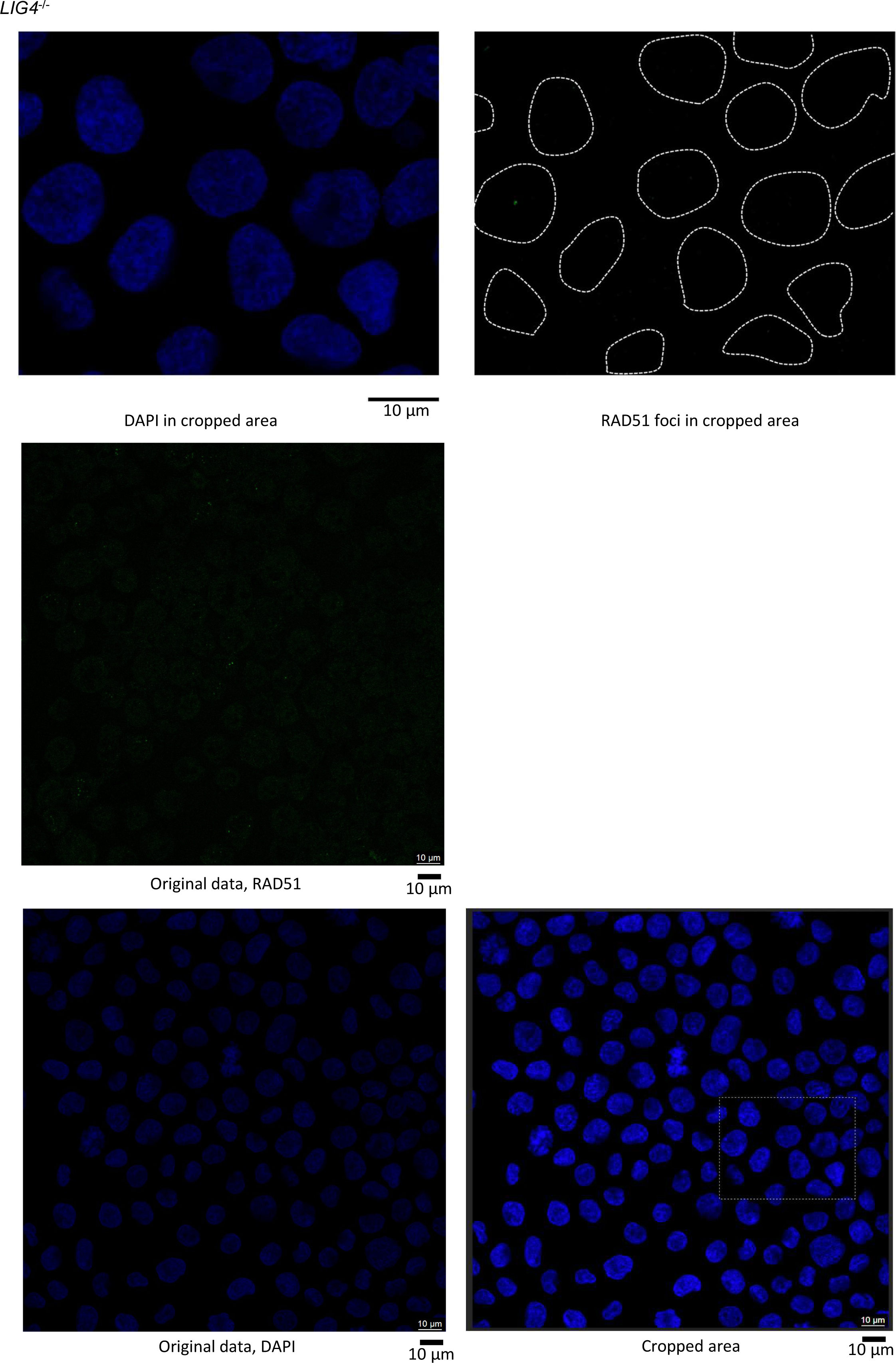
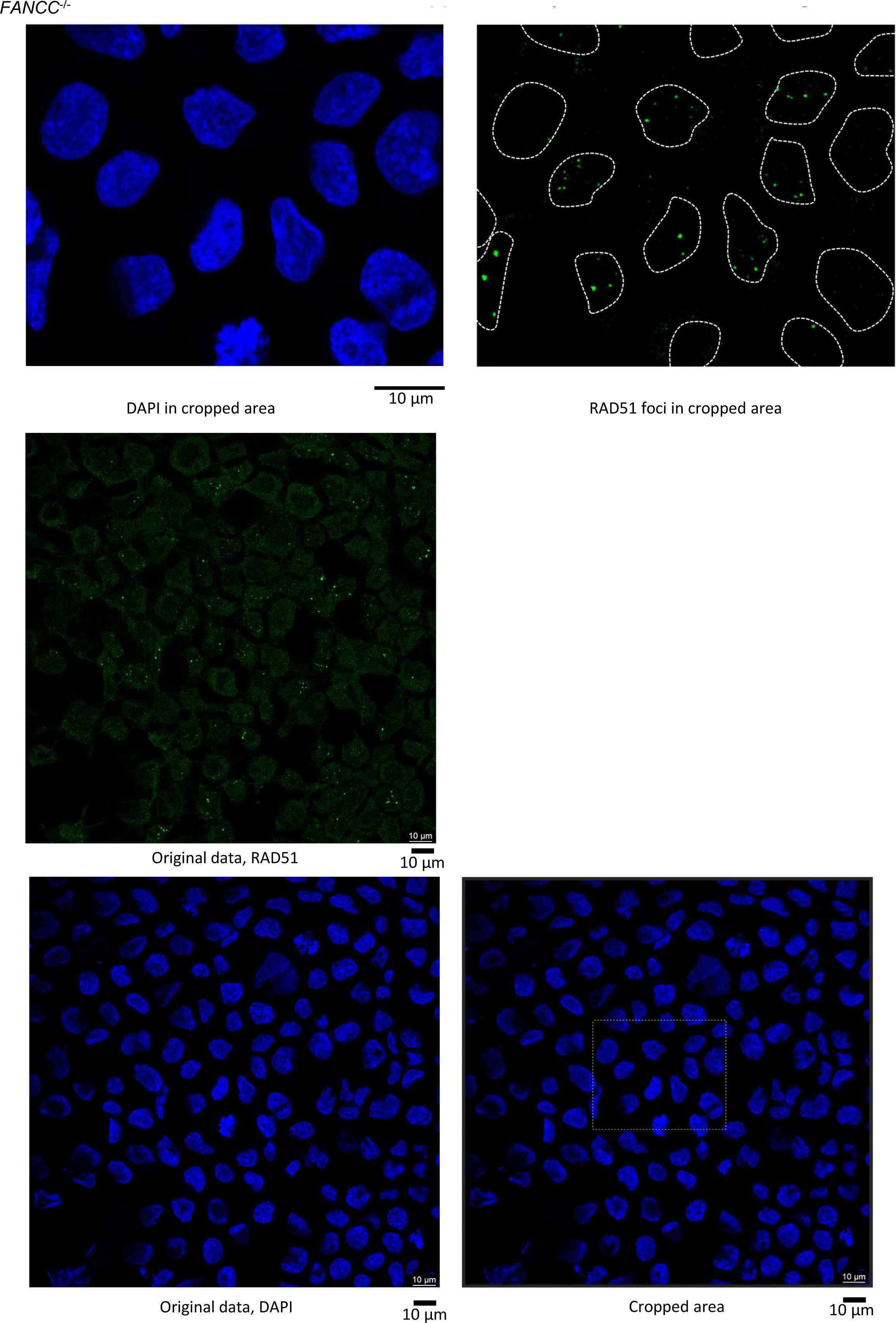
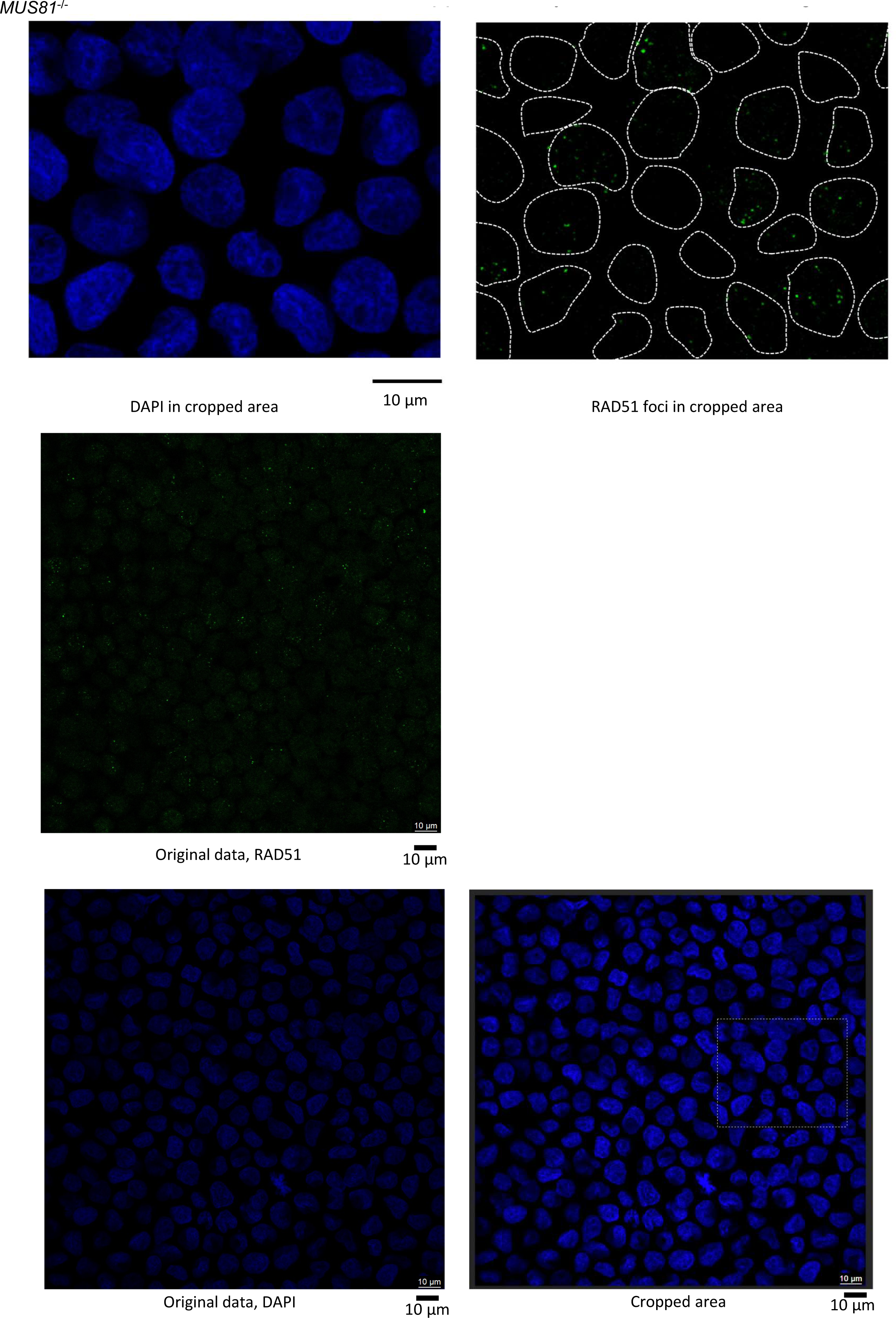
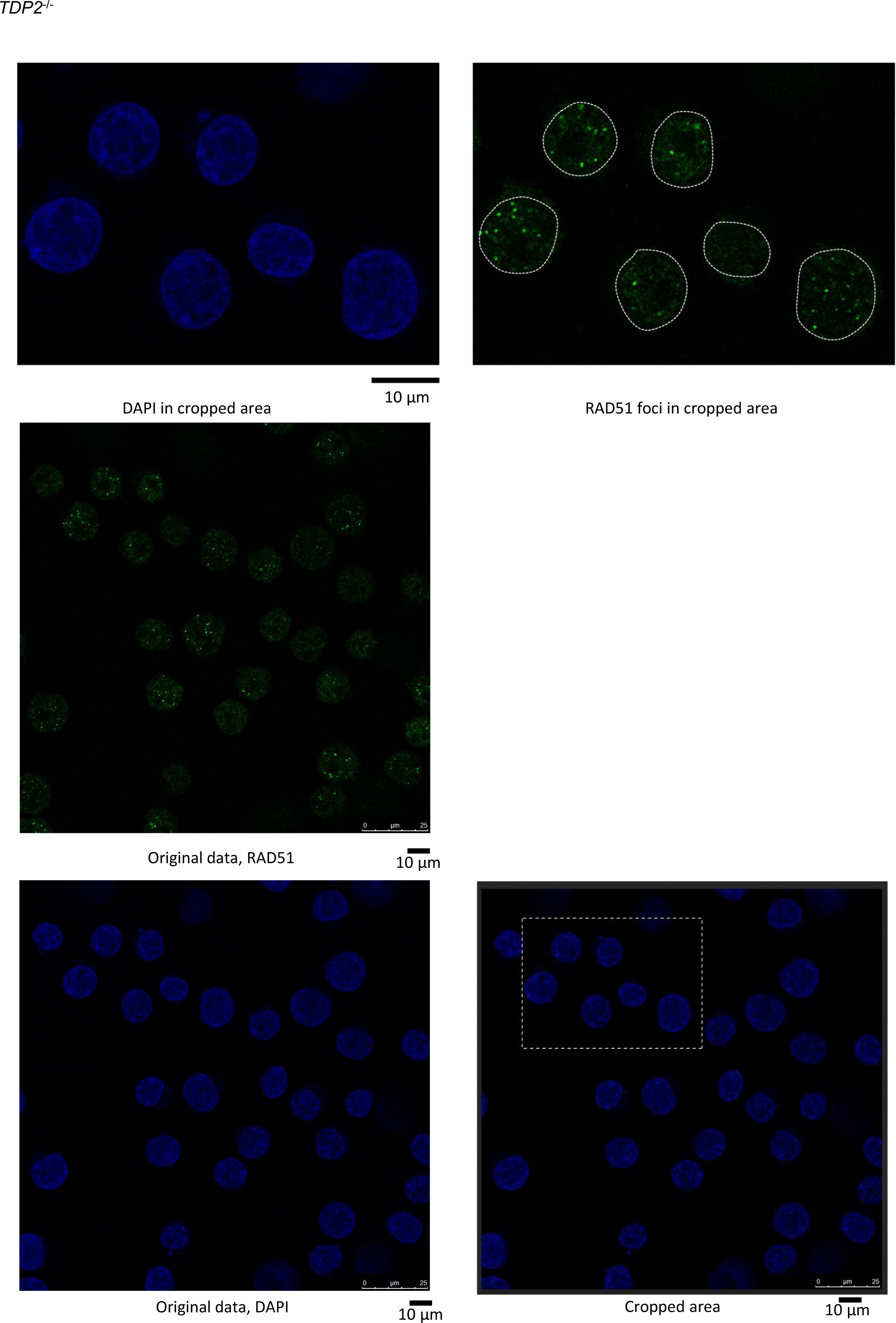
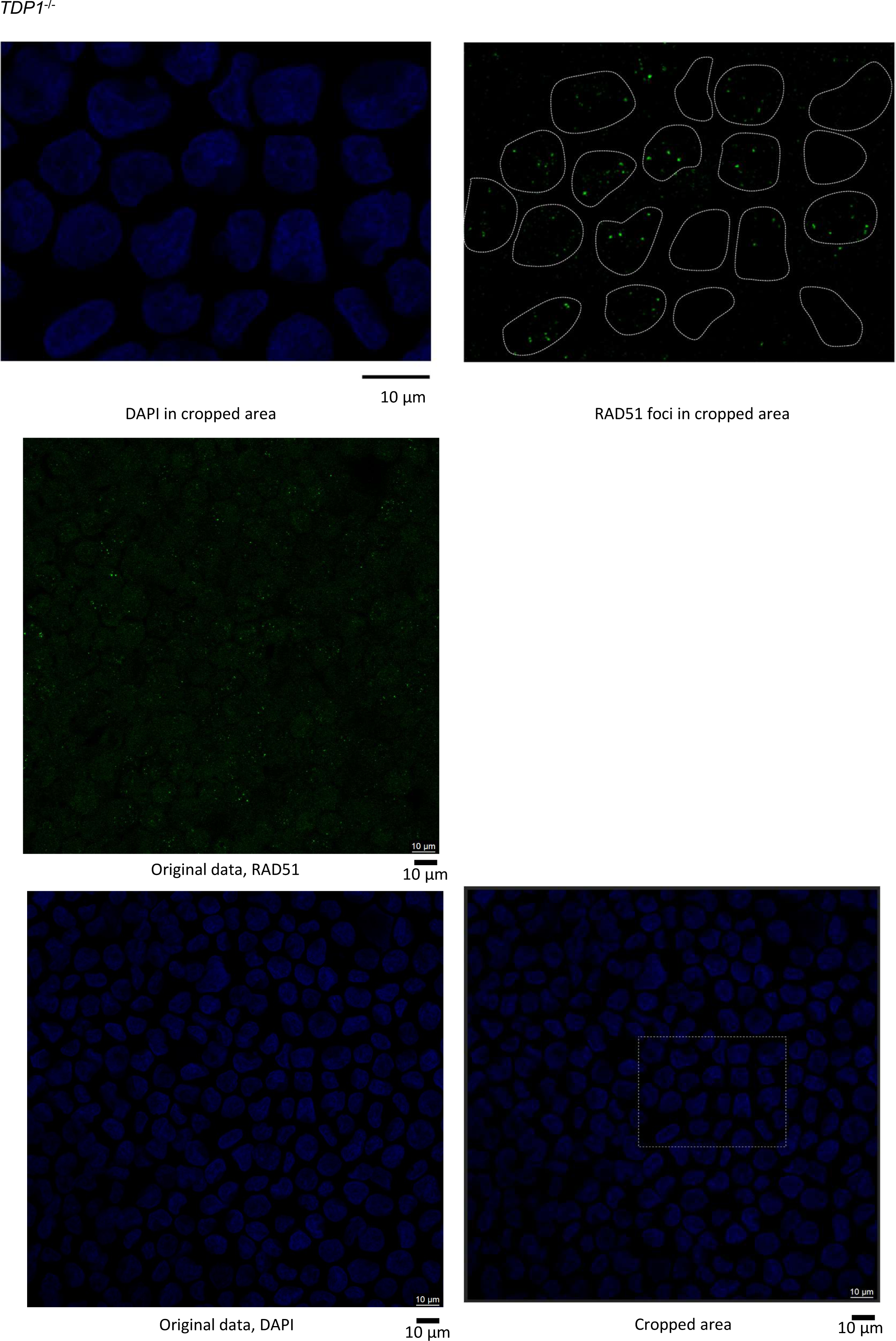

**Figure S3.**
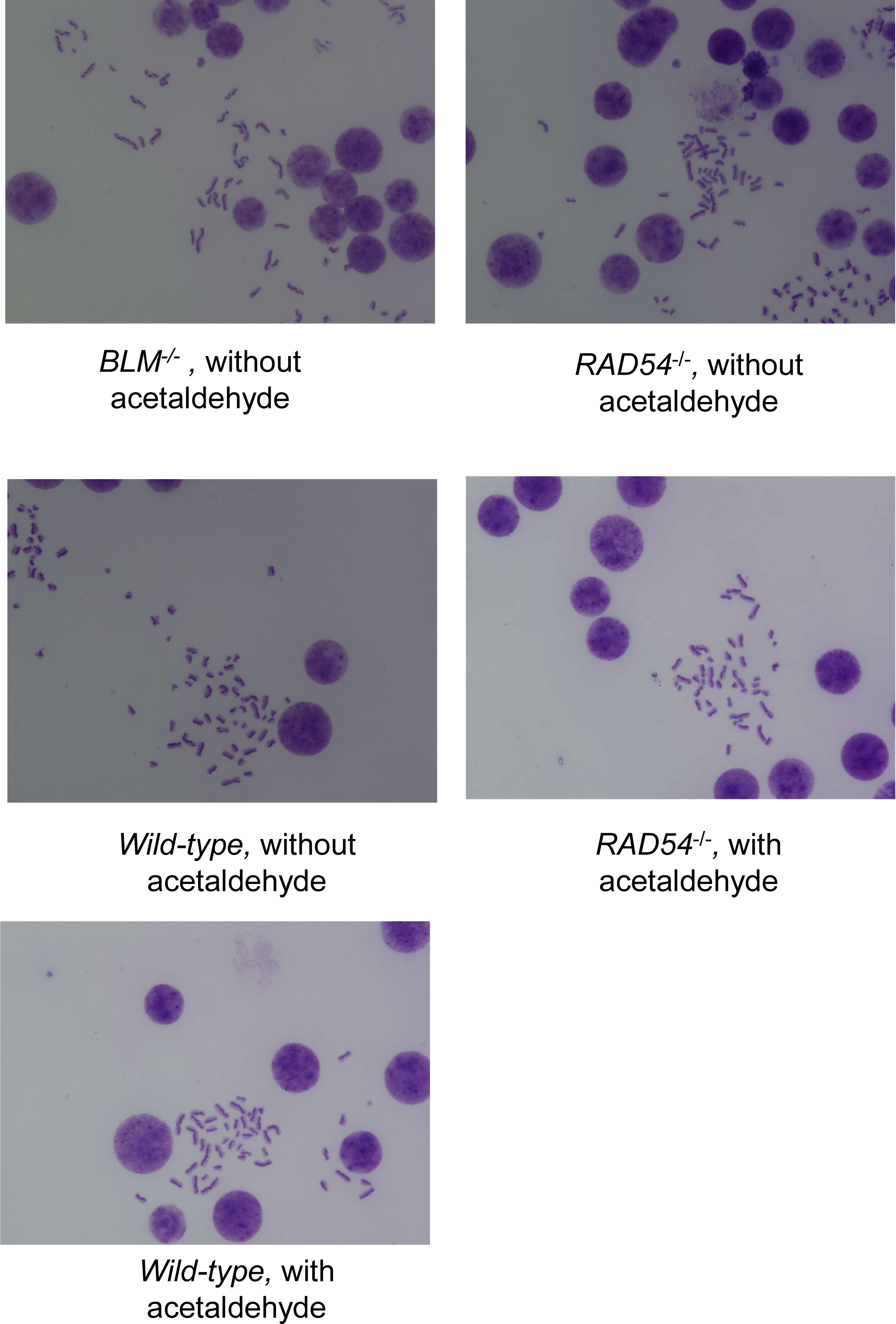

